# Synthetic STING agonists elicit powerful vaccine adjuvancy providing robust central memory and anti-tumour effects

**DOI:** 10.1101/2023.01.04.522614

**Authors:** Laurence D. Towner, Lekh N. Dahal, Martin C. Taylor, Kerry L. Cox, Tatyana Inzhelevskaya, Matthias Mack, Stephen R Wedge, Caroline Richardson, Mark S. Cragg, Stephen A. Beers

**Affiliations:** Antibody and Vaccine Group, Centre for Cancer Immunology, School of Cancer Sciences, Faculty of Medicine, University of Southampton, Tremona Road, Southampton SO16 6YD, UK; Centre for Drug Safety Science, Department of Pharmacology & Therapeutics, University of Liverpool, UK; Department of Nephrology, University Hospital Regensburg, 93053 Regensburg, DE; Cancer Research UK Newcastle Drug Discovery Unit, Clinical and Translational Research Institute, Newcastle University, Newcastle-upon-Tyne, UK; Astex Pharmaceuticals, Cambridge, UK

**Keywords:** CD8 T cells, STING, Type I IFN, vaccine, adjuvant, T cell memory, STING agonist

## Abstract

Drugs that target the innate immune sensor STING are known to be effective in modulating the immune infiltrate of the tumour microenvironment. STING agonists have potential to enhance responses to checkpoint inhibitor therapy, however, their ability to influence and shape adaptive immune responses is poorly understood. Here, we investigated the impact of a range of synthetic STING agonists on antigen specific CD8^+^ T-cell responses to soluble antigen using the murine OT-1 adoptive transfer model with Ovalbumin as the antigen to monitor T cell responses. Our data demonstrate that synthetic STING agonists are able to stimulate antigen specific T-cell expansion in response to challenge in mice. This effect required expression of STING, an intact myeloid compartment and Type-I IFN and TNFα signalling. Expanded T-cells post treatment differed from those induced by the established immune adjuvant, anti-CD40 antibody through lower induction of the immune checkpoint receptor PD-1. Furthermore, our data revealed a marked increase in the induction and persistence of CD8^+^ central memory cells after STING agonist and antigen challenge. Finally, we demonstrate that following rechallenge, STING agonism produced larger secondary responses that could be translated into enhanced tumour protection and survival. Therefore, synthetic STING agonists are capable of acting as potent immune adjuvants and can induce robust memory formation leading to better recall and tumour control. Critically, these benefits along with the lower expression of PD-1, have implications for their use as adjuvants for multiple immunotherapy and vaccine applications.

## Introduction

Innate immunity can be stimulated by various pattern recognition receptors (PRR) in response to pathogens or host cell components released during cell death or damage.[1] Stimulator of Interferon Genes (STING) is an endoplasmic reticulum (ER) resident intracellular PRR known to induce Type-I Interferons (IFN) in response to viral infection.[2, 3] STING activation occurs upon binding of cyclic dinucleotides (CDNs), either released by bacteria or produced by Cyclic GMP-AMP Synthase (cGAS) in response to cytoplasmic DNA.[4] Binding of CDNs leads to STING oligomerisation at the ER membrane and trafficking to the Golgi complex allowing the binding and sequential phosphorylation of TANK binding kinase 1 (TBK1) and interferon regulatory factor 3 (IRF3) with activated IRF3 then translocating to the nucleus to drive the transcription of interferon-responsive genes.[3, 5-8] Evidence suggests that STING induced Type-I IFN production is integral to adaptive T cell responses against tumour antigens through recruitment and promotion of antigen presenting cells and subsequent priming and expansion of CD8^+^ T cells.[9-11] For example, in preclinical models of syngeneic transplantable tumours, mice deficient in STING, IFR3 or Type-I IFN receptor (IFNAR) show defective T cell priming against tumour antigens and failure of immunogenic tumour rejection.[10, 11] A role for tumour necrosis factor alpha (TNFα) has also been described in T cell responses to STING activation in vivo.[12] Although antigen presenting cells, particularly dendritic cells (DC) are deemed indispensable for STING mediated antigen-specific T cell priming, molecular mechanisms underpinning STING activation in other immune cell subsets, and the nature and dynamics of the antigen primed T cell response is underexplored.[13] This has direct relevance to intrinsic tumour immunity in STING expressing tumour microenvironments (TME),[10] opening the possibility of therapeutic intervention to overcome immunosuppression within the TME.[14, 15] Recent studies, including ours,[16] have shown that stimulating this pathway has powerful immunostimulatory and anti-cancer effects, primarily mediated by macrophages. We have shown that STING agonists (STINGa) can overcome immunosuppression in the TME, reversing tumour associated macrophage inhibitory profiles to produce strong adjuvant effects for antibody immunotherapy. Importantly, primary human macrophage responses mirror those of murine macrophages, supporting the translational potential of these findings.[16] Similarly, others have shown that STING activation can improve response to checkpoint blockade therapy.[17-20] Currently 7 different synthetic STINGa, are being investigated in 10 clinical development programmes aimed at the treatment of solid tumours,[21] the majority in combination with checkpoint blockade (Table 1). Successful application of STING activation in the context of different immunomodulatory co-treatments will be reliant on a full understanding of the immune effects triggered by STING activation. Furthermore, understanding the determinants that govern optimal STING agonism and immune adjuvancy is urgently needed to contextualise their clinical performance and inform the requirements for success.

**Table 1.** Synthetic STING agonist in clinical development. Active clinical development programs involving synthetic STING agonist. Data sourced from ClinicalTrials.gov as of the index date of April 20, 2021.

| Compound | Co-therapy | NCT Number | Status | Conditions | Phases | Sponsor/Collaborators |
| --- | --- | --- | --- | --- | --- | --- |
| ADU-S100 | None | NCT03937141 | Active, not recruiting | Metastatic Head and Neck Cancer/Recurrent Head and Neck Cancer | II | Chinook Therapeutics, Inc. |
|  | Ipilimumab | NCT02675439 | Active, not recruiting | Advanced/Metastatic Solid Tumors or Lymphomas | I | Aduro Biotech, Inc./Novartis Pharmaceuticals |
| BI 1387446 | Ezabenlimab | NCT04147234 | Recruiting | Advanced/Metastatic Solid Tumors | I | Boehringer Ingelheim |
| BMS-986301 | Nivolumab/Ipilimumab | NCT03956680 | Recruiting | Advanced Solid Cancers | I | Bristol-Myers Squibb |
| CDK-002 | None | NCT04592484 | Recruiting | Advanced Solid Tumor | I | Codiak BioSciences |
| E7766 | None | NCT04144140 | Recruiting | Lymphoma/Advanced Solid Tumors | I | Eisai Inc./H3 Biomedicine Inc. |
|  | None | NCT04109092 | Withdrawn | Urinary Bladder Neoplasms | I | Eisai Inc./H3 Biomedicine Inc. |
| GSK3745417 | Pembrolizumab | NCT03843359 | Recruiting | Neoplasms | I | GlaxoSmithKline |
| MK-1454 | Pembrolizumab | NCT04220866 | Active, not recruiting | Head and Neck Squamous Cell Carcinoma (HNSCC) | II | Merck Sharp & Dohme Corp. |
|  | Pembrolizumab | NCT03010176 | Recruiting | Solid Tumors/Lymphoma | I | Merck Sharp & Dohme Corp. |
| MK-2118 | Pembrolizumab | NCT03249792 | Recruiting | Solid Tumor/Lymphoma | I | Merck Sharp & Dohme Corp. |
| SB 11285 | Atezolizumab | NCT04096638 | Recruiting | Melanoma/Head and Neck Squamous Cell Carcinoma/Solid Tumor | I | F-star Therapeutics, Inc. |
| SNX281 | None | NCT04609579 | Recruiting | Advanced Solid Tumors and Lymphoma | I | Stingthera, Inc. |
| SYNB1891 | Atezolizumab | NCT04167137 | Recruiting | Metastatic Solid Neoplasm/Lymphoma | I | Synlogic/IQVIA Biotech |
| TAK-676 | Pembrolizumab | NCT04420884 | Recruiting | Advanced Solid Cancers | I | Takeda |
|  | Pembrolizumab/Image-guided radiation therapy | NCT04879849 | Not yet recruiting | Carcinoma, Non-Small-Cell Lung/Triple Negative Breast Neoplasms/Squamous Cell Carcinoma of Head and Neck | I | Takeda |

Currently there is a growing interest in PRR ligands as vaccine adjuvants to prime innate immunity and boost antigen-specific T cell responses.[22] Vaccine models offer an ideal opportunity for understanding STINGa influence on T cell immunity and to dissect the key characteristics required for optimal responses. The capacity of 2’3’ cGAMP, the natural metazoan ligand for STING, to act as an adjuvant for optimal T cell responses in a vaccine setting has previously been described[23] but synthetic STINGa are underexplored in this context. Here we provide an extensive characterisation of the nature and dynamics of the T cell response in vaccine and anti-cancer settings following STING pathway activation via 2 synthetic STINGa: 5,6-dimethylxanthenone-4-acetic acid (DMXAA) a non-clinical murine specific agonist;[24-26] and a dithio-substituted diastereomer CDN with mixed linkage, ADU-S100, a clinically relevant agonist. We explore their activity as immune adjuvants and their capacity to support memory responses that confer tumour protection.

## Methods

### Mice, OT-I Adoptive transfer and tumour challenge

Wild-type C57BL/6, STING KO (*MPYS*^*-/-*^, generously provided by John Cambier[27]) and OT-I transgenic mice,[28] were bred and maintained in-house within the University of Southampton Biomedical Research Facility: specific pathogen free conditions, food (irradiated) and water was available ad-lib, mice were given 12 hour light/dark cycle and environmental enrichment; temperature maintained 20-24°C. Genotypes confirmed by PCR/flow cytometry. 10^4^ naïve OT-I splenocytes were delivered i.v. to WT C57BL/6 recipient mice (8-12 weeks, female) on d0. On d1, mice were challenged with 5 mg ovalbumin (Ova)(Sigma) i.p. alongside 100 μg mIgG1 anti-CD40 mAb (3/23, mouse IgG1 generated in-house as described previously[29, 30]) and the indicated doses of DMXAA(Invivogen), ADU-S100 was obtained from Oxeltis(Montpellier, France) and spin filtered using Amicon Ultra4 centrifugal filter tubes (Merck-Millipore, Watford, UK). Absence of endotoxins (<0.5 EU/mg) was confirmed by Wickham Laboratories (Gosport UK). STINGa dosed animals received a second dose on d2. For receptor blockade/cellular depletion experiments animals received mAbs by i.p. on d-1, and 0 of adoptive transfer: 500 µg anti-CSF1R (AFS98) further dosing on d1; 500 µg anti-CSF1 (5A1) further dosing d1&2; 500 µg anti-TNFalpha (XT3.1) further dosing on d1 (all preceding from BioXCell); 500 µg anti-IFNAR (MAR1-5A1) further dosing on day 1; 500 µg anti-Ly6G (1A8)(preceeding from Leinco); 75 µg anti-CCR2 (MC21)(further dosing days 1 and 2).[31] In some experiments mice were re-challenged with 30 nM Ova_257–264_ peptide i.v.. In other experiments mice were challenged s.c. with the Ova expressing, E.G7-Ova (ATCC)[32] and tumour growth monitored. Mice were culled once the humane endpoint had been reached (size 15×15 mm).

### Flow Cytometry

The following reagents were used for flow cytometry staining: (i) PE-labelled H-2Kb/SIINFEKL tetramer (in house), (ii) CD8a-APC (53-6.7), CD44-FITC (IM7), Ly6C-PerCPCy5.5 (HK1.4), Ly6G-PECy7 (RB6-8C5), CD11b-PE (M1/70)(preceding from eBiosciences), (iii) CD62L-PB (MEL-14), PD-1-PECy7 (RMP1-30)(preceding from Biolegend), (iv) F480-PB (CI:A3-1)(BioRad), (v) anti-mSTING (41)(Sigma Aldrich) and (vi) anti-rtIgM-AF647 (Invitrogen). For surface markers, cells were stained for 30m on ice in 1:1 PBS for blood or in PBS supplemented with 10% BSA and 10 mM NaN_3_ (FACswash) for splenocytes. Cells were then fixed in 1x erythrolyse buffer (BioRad) and washed in FACSwash. For intracellular markers, cells were stained using FoxP3 Intracellular staining kit, (eBiosciences). Stained cells were acquired on a FACsCanto II (BD Biosciences) and data analysed using FlowJo v10 (FlowJo LLC).

### T cell proliferation

Total T cells were purified from mouse splenocytes using an EasySep murine T cell isolation kit and labelled with 5 μM CFSE. 100,000 purified T cells were plated in a 96 well UB plate (100 μl/well). Anti-CD3/anti-CD28 coated dynabeads were used to stimulate the cells with/without STINGa for 72 hours. Cells were then stained with anti-CD8-APC and analysed by flow cytometry.

### Statistics

Statistics were calculated using Graphpad Prism software. To compare 3 or more groups an unpaired, one-way ANOVA with Tukey’s adjustment was performed, or for 2 group comparisons an unpaired, two-tailed Students t-test was used. Where applicable, AUC was calculated and pertinent statistical test performed. Kaplan–Meier curves were produced and analyzed by Log rank (Mantel– Cox) test. Significance of comparisons are indicated on plots: ns = not significant; * p<0.05; ** p< 0.01; *** p< 0.001; **** <0.0001.

## Results

### STING agonists are potent immune adjuvants, inducing robust antigen specific CD8 T cell responses

We first investigated the ability of the murine specific STINGa DMXAA to stimulate antigen specific CD8 T cell (CD8^+^) response to the model antigen Ovalbumin (Ova) and compared it with an established immune adjuvant, the agonistic anti-CD40 antibody, 3/23.[28, 33-35] Ova-specific OT-I splenocytes were transferred into C57BL/6 wild type mice and 3 days later recipients were challenged with Ova along with maximally stimulating doses of anti-CD40 (for dose-response see Supplementary Figure 1A) or DMXAA, and peripheral Ova specific CD8^+^ responses monitored over time using SIINFEKL tetramer staining and flow cytometry (Figure 1A). Ova alone did not induce a response, however, co-administration with either anti-CD40 or DMXAA induced marked Ova-specific CD8^+^ (CD8^+^Tet^+^) expansion that peaked 7 days post-challenge, accounting for 40-80% of all peripheral blood leukocytes (singlets) and 80-90% of CD8^+^s (Figure 1B & C, respectively). Notably, DMXAA induced larger and longer lasting responses as compared to anti-CD40 with CD8^+^Tet^+^ cells typically expanding to ∼75% singlets and ∼90% of CD8^+^s at their peak and remaining at ∼70% of CD8^+^s 7 weeks later (day 49). DMXAA adjuvancy was dose dependent up to and including the maximal tolerated dose of 400 μg with 100 and 200 μg doses having a more modest impact on expansion as a proportion of both singlets and CD8^+^s (Figure 1D). We also confirmed that the observed CD8^+^Tet^+^ expansion in response to DMXAA was STING dependent, as the DMXAA agonist effect was completely abolished in STING knockout (KO) mice whilst anti-CD40 retained its activity (Figure 1E). Given the STING dependence of the DMXAA adjuvant effects observed, we next investigated the contribution of Type-I IFN and TNFα, both demonstrated previously to be important for STING induced CD8^+^ responses. In keeping with reports from tumour models[15, 36] blockade of either IFNAR or TNFα largely abrogated the DMXAA induced responses (Figure 1F), further supporting their importance in a vaccination setting. To confirm the broader applicability of the above observations we then assessed the clinically relevant STINGa ADU-S100[37]. In comparison to 2’3’ cGAMP, DMXAA is 2-fold less potent and ADU-S100 4-fold more potent,[37, 38] giving a theoretical potency difference between DMXAA and ADU-S100 of ∼8-fold. ADU-S100 induced CD8^+^Tet^+^ expansion with kinetics similar to anti-CD40 and with similar dose-dependence to DMXAA (Figure 1 G & H). These data support the contention that synthetic STINGa are potent adjuvants for antigen specific CD8^+^ vaccine responses and that these effects are STING, Type-I IFN and TNFα dependent.

**Figure 1.**
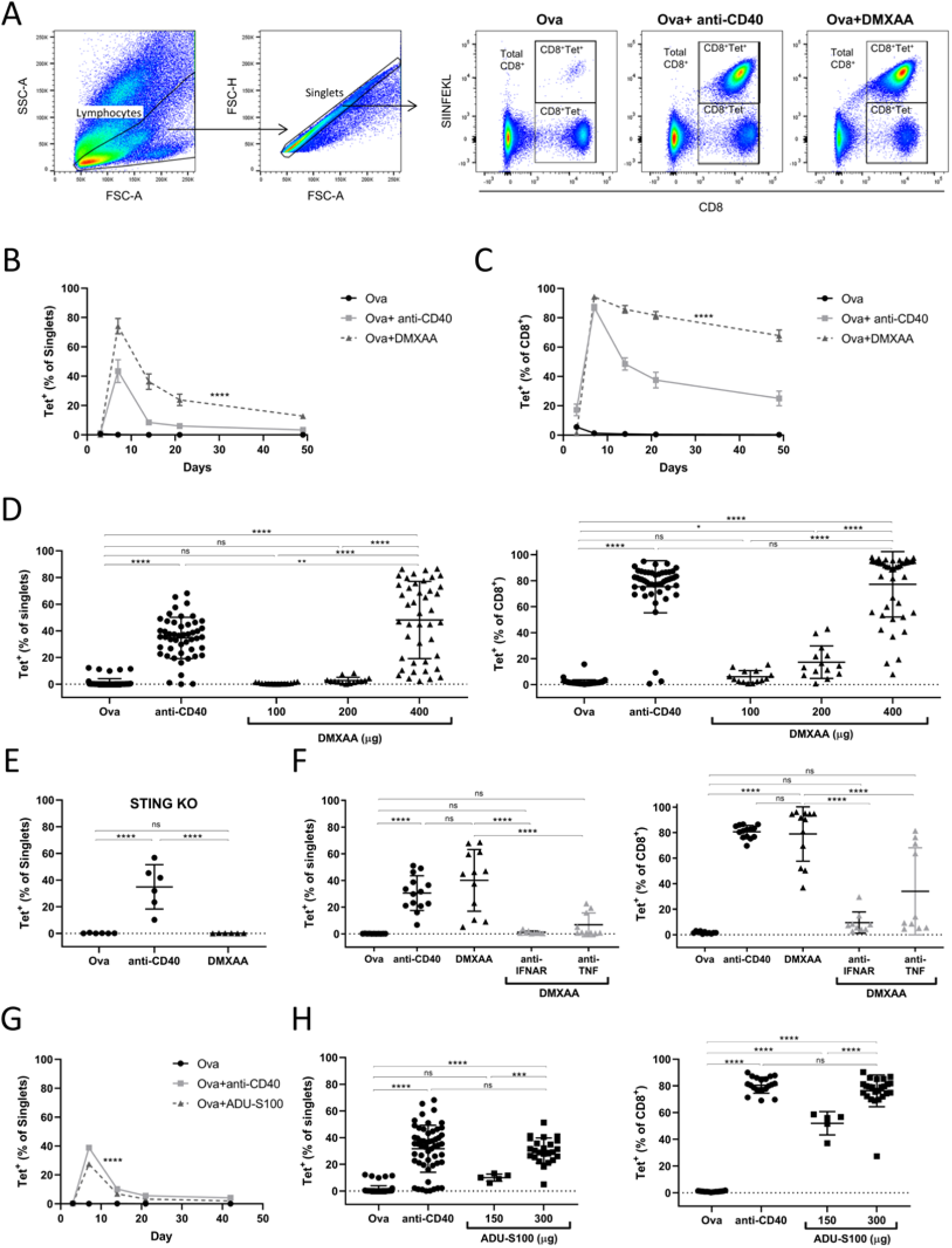
Synthetic STING agonist are potent adjuvants in a vaccination setting and induce robust antigen specific CD8 T cell expansion dependent upon type I IFN and TNFα. WT C57BL/6 mice received adoptive transfer of OT-I splenocytes and challenged with 5 mg Ovalbumin (Ova) and treated with either DMXAA, anti-CD40 or carrier control and CD8^+^ T cells monitored in blood. (**A**) Representative flow cytometry plots showing detection of CD8^+^Tet^+^ and CD8^+^Tet^-^ cells present in blood on day 7 post challenge. (**B**&**C**) Representative CD8^+^Tet^+^ expansion kinetics, shown as both %Singlets (**B**) and %CD8s (**C**). (**D**) CD8^+^Tet^+^ expansion at peak (day 7); presented as %Singlets (left) and %CD8s (right) (n=≥14). (**E**) Peak CD8^+^Tet^+^ expansion as %Singlets in STING KO mice (n=6). (**F**) Peak CD8^+^Tet^+^ expansion in mice receiving DMXAA with IFNAR and TNF blockade; presented as %Singlets (left) and %CD8s (right) (n=≥9). (**G**) CD8^+^Tet^+^ expansion kinetics after ADU-S100 administration. (**H**) Peak CD8^+^Tet^+^ in mice receiving ADU-S100 presented as %Singlets (left) and %CD8s (right) (n=≥5). Each point represents an individual mouse, mean is indicated with error bars showing SD. *P* values were determined using One-way ANOVA with Tukey’s adjustment/ two-tailed Students t-test. ns = not significant; * *P*<0.05; ** *P*< 0.01; *** *P*< 0.001; \*\*\*\**P*<0.0001.

### STING agonists elicit more restricted PD-1 expression on expanded antigen specific CD8^+^s compared to anti-CD40 antibody

The ability of synthetic STINGa to induce rapid and robust T cell proliferation in the context of a soluble antigen, and in some cases produce a greater expansion than other established adjuvants (anti-CD40), led us to investigate the mechanisms that regulate these T cell responses. Programme cell death receptor 1 (PD-1) is upregulated upon TCR engagement and limits further T cell activation upon engagement with PD-1 ligands.[39, 40] Expression of PD-1 on CD8^+^s (total, Tet^+^ and Tet^-^), was monitored at the peak of expansion post-challenge (day 7) (Figure 2A). Administration of either anti-CD40 or DMXAA alongside Ova induced significantly more PD-1 expression compared to Ova alone (Figure 2B) in all CD8^+^ populations, but DMXAA induced significantly less PD-1 than anti-CD40 (Figure 2B). These analyses demonstrate reduced PD-1 induction on expanded cells after DMXAA treatment (CD8^+^Tet^+^ and CD8^+^Tet^-^) compared to anti-CD40, despite the overall expansion being greater with DMXAA. To confirm these findings, dose titrations were carried out and fold change in levels of PD-1 expression on CD8^+^s following anti-CD40 or DMXAA administration compared to Ova alone plotted. Overall, ∼4-fold upregulation of PD-1 was induced at the peak of the response by anti-CD40. It was significantly lower for all doses of DMXAA tested, up to the maximal tolerated dose where it reached ∼2.5-fold. Lower doses of DMXAA induced little to no PD-1 (Figure 2C) reflecting the muted expansion of CD8^+^s as observed in Figure 1D. In contrast, anti-CD40 induced similar levels of PD-1 on CD8^+^Tet^+^ populations across all doses tested (Supplementary Figure 1B). To confirm that the observed differences were characteristic of STING pathway activation, rather than the particular murine-specific STINGa, we performed similar analysis with ADU-S100 (Figure 2D). We observed that the highest dose of ADU-S100, like DMXAA, induced levels of PD-1 that were significantly lower than those induced by anti-CD40. Notably, although lower than anti-CD40, ADU-S100 induced significantly higher levels of PD-1 compared to DMXAA at comparable doses (Supplementary Figure 1C). Furthermore, the upregulation of PD-1 was STING pathway activation dependent as it was completely abrogated in mice deficient for STING (Figure 2E) or in mice that received blocking antibodies against TNF-α or Type-I IFN (Figure 2F), mirroring the curtailed CD8^+^ responses. These data indicate that synthetic STINGa induce a CD8^+^ response during vaccination, which is characterised by restricted PD-1 induction on expanded cells compared to anti-CD40, but also highlight that the nature of the response to different synthetic STINGa can vary, suggesting that it may be possible to optimise such reagents for combination with immunotherapy.

**Figure 2.**
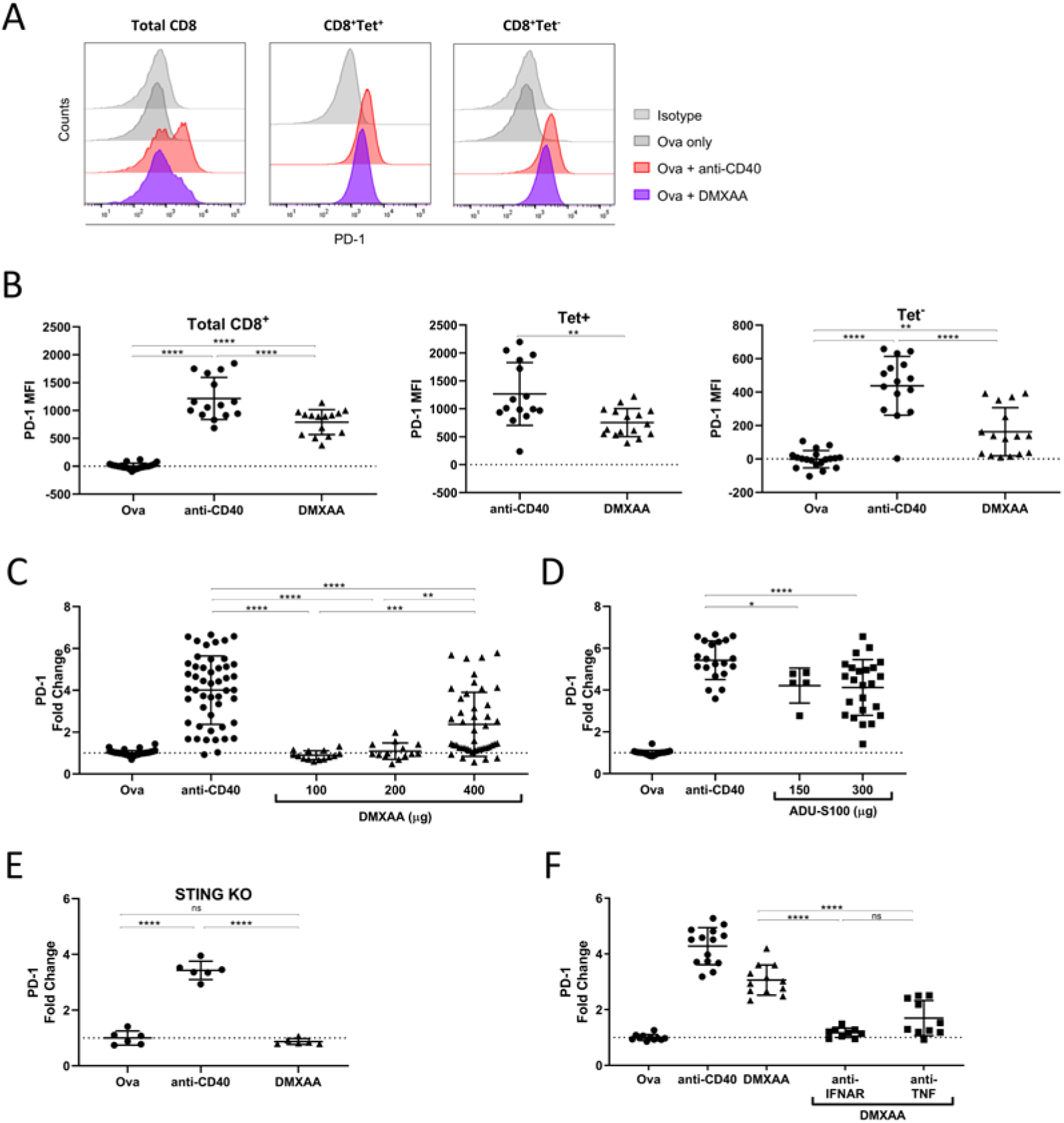
PD-1 expression on expanded antigen specific CD8 T cells is lower when induced by STING agonist compared to anti-CD40. (**A**) Representative flow cytometry plots showing PD-1 expression on total CD8, CD8^+^Tet^+^and CD8^+^Tet^-^ populations in mice treated with anti-CD40 and DMXAA compared to Ova alone and isotype control. (**B**) PD-1 expression for total CD8s, CD8^+^Tet^+^ and CD8^+^Tet^-^ (n=≥15) (**C**-**F**) PD-1 expression on total CD8s as a fold difference compared to Ova challenge alone: (**C**) DMXAA dose titration and (n=≥14), (**D**) ADU-S100 dose titration (n=≥5), (**E**) DMXAA in STING KO mice, and (**F**) DMXAA with blockade of IFNAR and TNF (n=≥9). Each point represents an individual mouse, mean is indicated with error bars showing SD. *P* values were determined using One-way ANOVA with Tukey’s adjustment. ns = not significant; * *P*<0.05; ** *P*< 0.01; *** *P*< 0.001; \*\*\*\**P*<0.0001.

### STING-induced vaccine responses demonstrate myeloid dependence

Next, we sought to confirm the cellular STING expression requirements for the STING agonist-mediated vaccine responses. We assessed STING expression levels in splenic immune cell populations from wild-type C57BL/6 mice by flow cytometry (Figure 3A). STING was found to be widely expressed in murine immune cells but was higher in lymphocytes with T cells>NK cells>monocytes and B cells. Myeloid cells expressed varying levels of STING with monocytes the highest followed by macrophages and with an absence of detectable STING in neutrophils. It is therefore possible that direct STING activation in T cells might account for, in part, the adjuvant effect observed. To test this, an in vitro proliferation assay was performed with isolated T cells stimulated with anti-CD3 in the presence of increasing concentrations of STINGa (Figure 3B). Both CD4^+^s and CD8^+^s showed a dose-dependent reduction in proliferation with a significant impact observed in the presence of 1 µg/mL, the EC_50_ value for DMXAA,[37] and complete abrogation at 10 µg/mL. These findings are concordant with the data in Figure 1E where a complete abrogation of STINGa-induced CD8^+^Tet^+^ expansion was observed when STING expressing OT-I T cells were adoptively transferred into STING KO recipient mice, supporting the contention that T cell intrinsic STING is insufficient to induce responses. Indeed, previous reports have indicated direct toxicity of STING activation in T cells which was shown to be mediated by stress and subsequent apoptosis.[41] These data suggest that direct agonism of STINGa on T cells cannot account for the marked adjuvant effect observed, instead indicating that sustained STING activation may serve to limit T cell expansion. Given our previous observations of the potent effects of STINGa on myeloid cells,[16] we sought to determine the role of myeloid cells in the STINGa-mediated vaccine responses. To assess this, the OT-I adoptive transfer model was repeated after first deleting/modulating different myeloid cell populations, such as neutrophils (anti-Ly6G), macrophages (anti-CSF1R and anti-CSF1) and monocytes (anti-CCR2)(Figure 3C). Results showed that targeting the CSF1 dependent myeloid cell population (largely macrophages and inflammatory monocytes) significantly reduced STING mediated adjuvancy as monitored by the expansion of CD8^+^Tet^+^ cells on day 7, whereas anti-CCR2 mediated depletion of monocytes showed a trend towards reduced expansion. Depletion of neutrophils had no effect on the T cell expansion. These data indicate that macrophages likely play an important role in mediating STING adjuvancy in this model.

**Figure 3.**
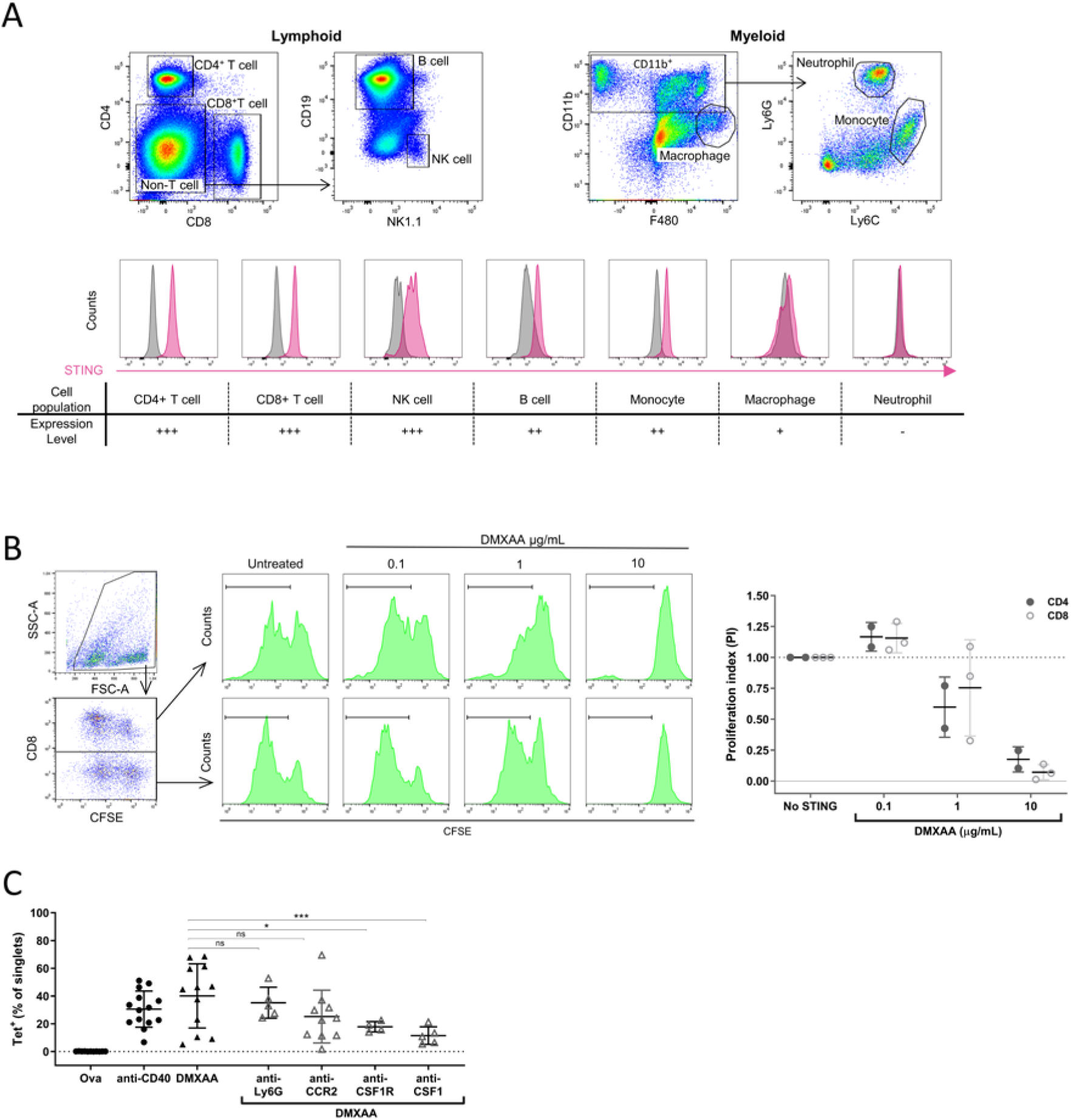
STING agonists require myeloid compartment for adjuvancy. (**A**) STING expression in mouse immune subsets: top panel; lymphoid and myeloid gating strategies, middle panel; plots comparing STING detection to isotype control, bottom panel; table summarising expression levels. (**B**) CD3/CD28 induced CD4^+^ and CD8^+^ T cell proliferation in the presence of DMXAA (n=3). (**C**) Day 7 CD8^+^Tet^+^ expansion induced by 400 µg DMXAA with concomitant immune cell depletions/receptor blockade: anti-Ly6G, neutrophils; anti-CSF1R, macrophage; anti-CSF1, CSF1 blockade; anti-CCR2 and CCR2 KO, monocytes (n=≥5). Each point represents an individual mouse/biological repeat, mean is indicated with error bars showing SD. *P* values were determined using One-way ANOVA with Tukey’s adjustment. ns = not significant; * *P*<0.05; ** *P*< 0.01; *** *P*< 0.001; \*\*\*\**P*<0.0001.

### STING agonists evoke a rapid, dominant and persistent central memory CD8 T cell population

*P P*

The low PD-1 response seen in Tet^+^ CD8s triggered by STINGa during immunisation with Ova was different to that evoked by anti-CD40, likely reflecting their respective induction of Type-I versus Type-II IFN.[3, 33] We therefore sought to understand whether these characteristics also led to an altered memory phenotype in the Ova-specific CD8^+^s induced. It is well established that upon priming, naïve CD8^+^s differentiate into effector cells which rapidly expand, then contract to leave a memory population that persists.[42, 43]

Using expression of CD62L and CD44 as markers we sought to define the relative proportions of Naive (CD62L^+^CD44^-^), effector/effector memory (E/EM; CD62L^-^CD44^+^) and central memory precursors/central memory (CMp/CM; CD62L^+^CD44^+^) cells in the CD8^+^ compartment post Ova challenge (Figure 4A and B). Figure 4B compares the kinetics of these various populations of Tet^+^ as a %Singlets (left) and %Tet^+^ (right) following maximal anti-CD40 or STINGa (DMXAA) treatment. Shown as %Singlets, the kinetics demonstrate peak expansion in both treatments to be dominated by E/EM but that CMp/CM induction is markedly greater after STINGa treatment. Contraction of E/EM was rapid with CMp/CM populations remaining stable after STINGa treatment compared to anti-CD40. This larger CMp/CM induction and its persistence led to an early crossover from EM to CM dominance in STINGa treated mice (∼ day 49 post-challenge) that did not occur with anti-CD40. Overall, these data indicate that STING agonism induces an increased proportion of CMp cells early in the response resulting in a greater proportion of memory cells 90 days after challenge compared to anti-CD40.

**Figure 4.**
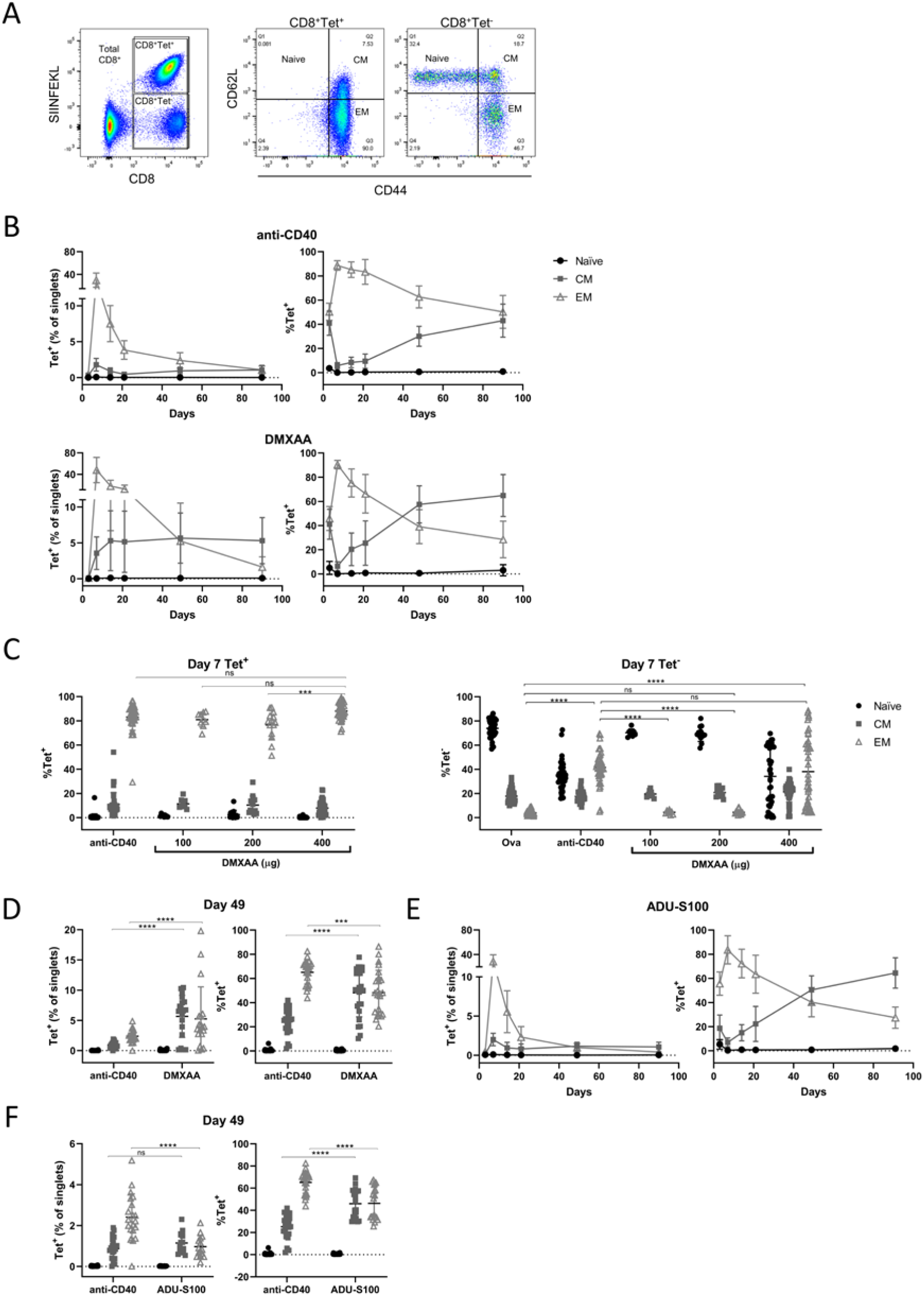
Antigen specific central memory CD8 T cells dominate memory responses in STING agonist treated mice. (**A**) Representative gating for defining memory status: naïve, CD62L^hi^CD44^lo^; central memory (CM), CD62L^hi^CD44^hi^; and effector memory (EM), CD62L^lo^CD44^hi^. (**B**) Memory phenotype kinetics of CD8^+^Tet^+^ cells post challenge with anti-CD40 (top) and DMXAA (bottom), presented as %Tet^+^ (left) and %Singlets (right) (n=≥15). (**C**) Memory status of Tet^+^ (left) and Tet^-^ (right) CD8s on day 7 in mice treated with DMXAA (n=≥9). (**D**) Memory response to DMXAA at day 49, presented as %Singlets (left) and %Tet^+^ (right) (n=≥14). (**E**&**F**) Memory phenotype of Tet^+^ cells in response to ADU-S100: (**E**) Kinetics; %Singlets (left) and %Tet^+^ (right), (**F**) memory phenotype on day 49 as %Singlets (left) and %Tet^+^ (right) (n=≥5). Each point represents an individual mouse, mean is indicated with error bars showing SD. *P* values were determined using One-way ANOVA with Tukey’s adjustment. ns = not significant; * *P*<0.05; ** *P*< 0.01; *** *P*< 0.001; \*\*\*\**P*<0.0001.

We subsequently explored the difference between anti-CD40 and STINGa further, looking at these same populations in Tet^+^ and Tet^-^ cells on day 7 (Figure 4C), and Tet^+^ on day 49 (Figure 4D) post-challenge with differing doses of DMXAA. On day 7, Tet^+^ cells were dominated by effector cells regardless of treatment or dosing (Figure 4C, left). In support of our earlier PD-1 observations, Tet^-^ cells showed a dominance of naïve cells in Ova alone control and low dose (100-200 μg) STINGa groups but a significant widening of the response, with some effector expansion, in anti-CD40 and high dose STINGa groups (Figure 4C, right). This widened response had disappeared by day 49 and 90 (Supplementary Figure 2A). On day 49, CM cells as %Singlets and %Tet^+^ were significantly increased in STINGa compared to anti-CD40 treated mice (Figure 4D). This confirmed that by day 49, Tet^+^ CM cells were beginning to dominate the memory pool in STINGa treated animals but not after anti-CD40 treatment. It remained unclear whether the different EM and CM kinetics between treatments was simply a reflection of differences in the magnitude of responses induced, with DMXAA frequently expanding CD8^+^s more potently than anti-CD40, or intrinsic to these different adjuvants.

To address this, we repeated these analyses with the clinically relevant STINGa, ADU-S100, which produced similar expansion magnitude and kinetics to anti-CD40 (Figure 1G). Here we observed that ADU-S100 induced Ova-specific CD8^+^ expansion (Figure 4E) with an early cross-over from E/EM to CMp/CM dominance similar to that observed with DMXAA and that this was due to greater CMp/CM persistence compared to anti-CD40. On day 49 E/EM populations as %Singlets in ADU-S100 treatment were significantly lower than in anti-CD40 treated mice (Figure 4F). The proportion of CM was greater though this did not reach significance. These observations translated to significantly more CMp/CM in ADU-S100 as %Tet^+^ and significantly less E/EM compared to anti-CD40. We also found similar patterns remained on day 90 in mice treated with both STINGa (Supplementary Figure 2B & C). This confirmed that both synthetic STINGa were capable of producing an early cross-over from EM to CM dominance resulting from a greater induction of CMp cells that persisted over the time course, suggesting a common characteristic of STINGa.

We then sought to understand the cellular dependence of these memory effects using the same cellular blockade and depletion approaches as before (Figure 3C). Figure 5A shows that where there was disruption of the initial expansion response, as with anti-CSF1 and to a lesser extent anti-CCR2, the switch from EM to CM dominance at day 49 was lost. This was confirmed by kinetics over 90 days post-challenge, which demonstrated that induction of CMp/CM was abrogated, translating into a loss of the cross-over effect (Figure 5B). These data support our previous assertion that macrophages, and to a lesser extent monocytes, are important for these memory effects with neutrophils being largely redundant.

**Figure 5.**
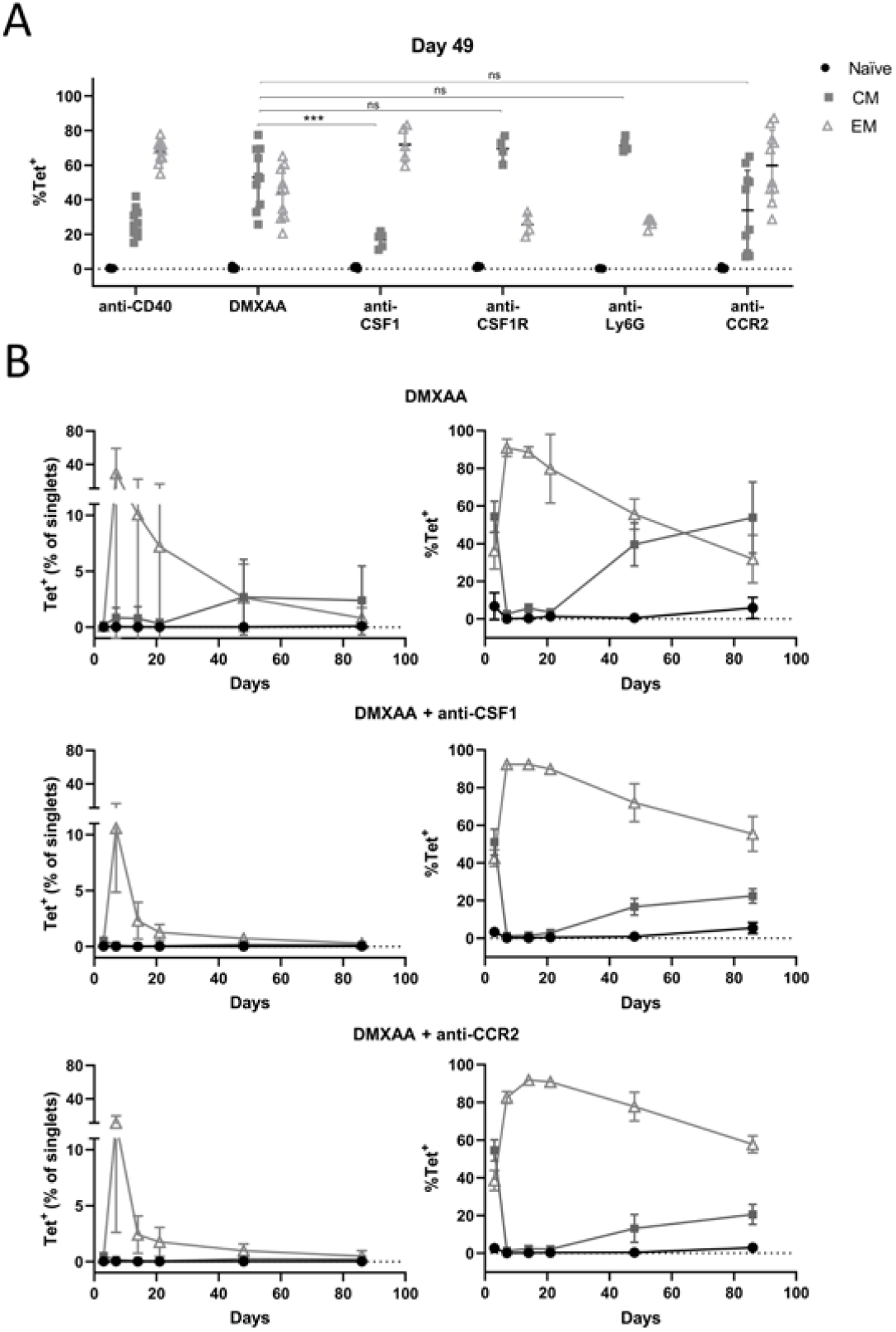
Depletion of monocytes and macrophages abrogates antigen specific central memory response with STING agonist. (**A**) Memory phenotype of Tet^+^ cells on day 49 induced by DMXAA with depletion of myeloid populations: neutrophils, anti-Ly6G; macrophages, CSF1R; CSF1 blockade, CSF1; and monocytes (n=≥4). (**B**) Memory phenotype kinetics of Tet^+^ cells as %Singlets (left) and % Tet^+^ (right) after (top panels) DMXAA, (middle) DMXAA+CSF1 blockade, and (bottom) and monocyte depletion (n=5). Each point represents an individual mouse, mean is indicated with error bars showing SD. *P* values were determined using One-way ANOVA with Tukey’s adjustment. ns = not significant; * *P*<0.05; ** *P*< 0.01; *** *P*< 0.001; \*\*\*\**P*<0.0001.

Together, these data show that STINGa, regardless of the magnitude of CD8 response, lead to a proportionately higher CMp response that persists leading to an earlier CM dominance when compared with anti-CD40 treatment.

### STING activation induced central memory correlates with effective anti-tumour immunity

We next investigated whether the robust CM response induced by STINGa translated into antigen-specific therapeutic benefit. First, we confirmed that antigen specific memory responses were generated and intact in mice immunised with Ova in the presence of STINGa (Figure 6A). The left panel shows that blockade of type-I IFN and TNFα inhibited the initial expansion of the primary response. Following re-challenge with the Ova derived SIINFEKL peptide, 90 days after initial Ova administration, we observed an expansion in CD8^+^Tet^+^ cells in mice previously co-treated with DMXAA and Ova, which peaked at ∼10% of total CD8s at day 7 whilst no response was evident in mice that received Type-I IFN or TNF-α blocking antibodies during primary challenge (Figure 6A, right panel). Next, immunised mice were challenged with Ova expressing E.G7 tumour cells, and monitored for tumour development and survival (Figure 6B & C). Despite the robust primary response to Ova 90 days earlier (Figure 6B, left panel) only 50% of the mice survived out to day 100 after anti-CD40 treatment whilst mice treated with 400μg DMXAA, received 100% protection against E.G7 tumour. The protection provided by DMXAA was dose-dependent with 200 μg providing better protection than 100 μg. Finally, we compared the tumour protection provided by ADU-S100 to that by DMXAA. Here, again DMXAA conferred complete protection from tumour challenge but in marked contrast, ADU-S100 showed a negative dose correlation to tumour protection with the highest dose evoking less tumour protection (Figure 6C). Taken together these data provide evidence that the robust CM response induced by STINGa provides long term, functionally relevant, memory and confers better tumour protection than anti-CD40. It is notable that, although DMXAA is ∼8 fold less potent than ADU-S100, the former provided superior tumour protection, suggesting that agonist potency may be a limiting factor in the generation of effective CM and the responses evoked by STING agonism.

**Figure 6.**
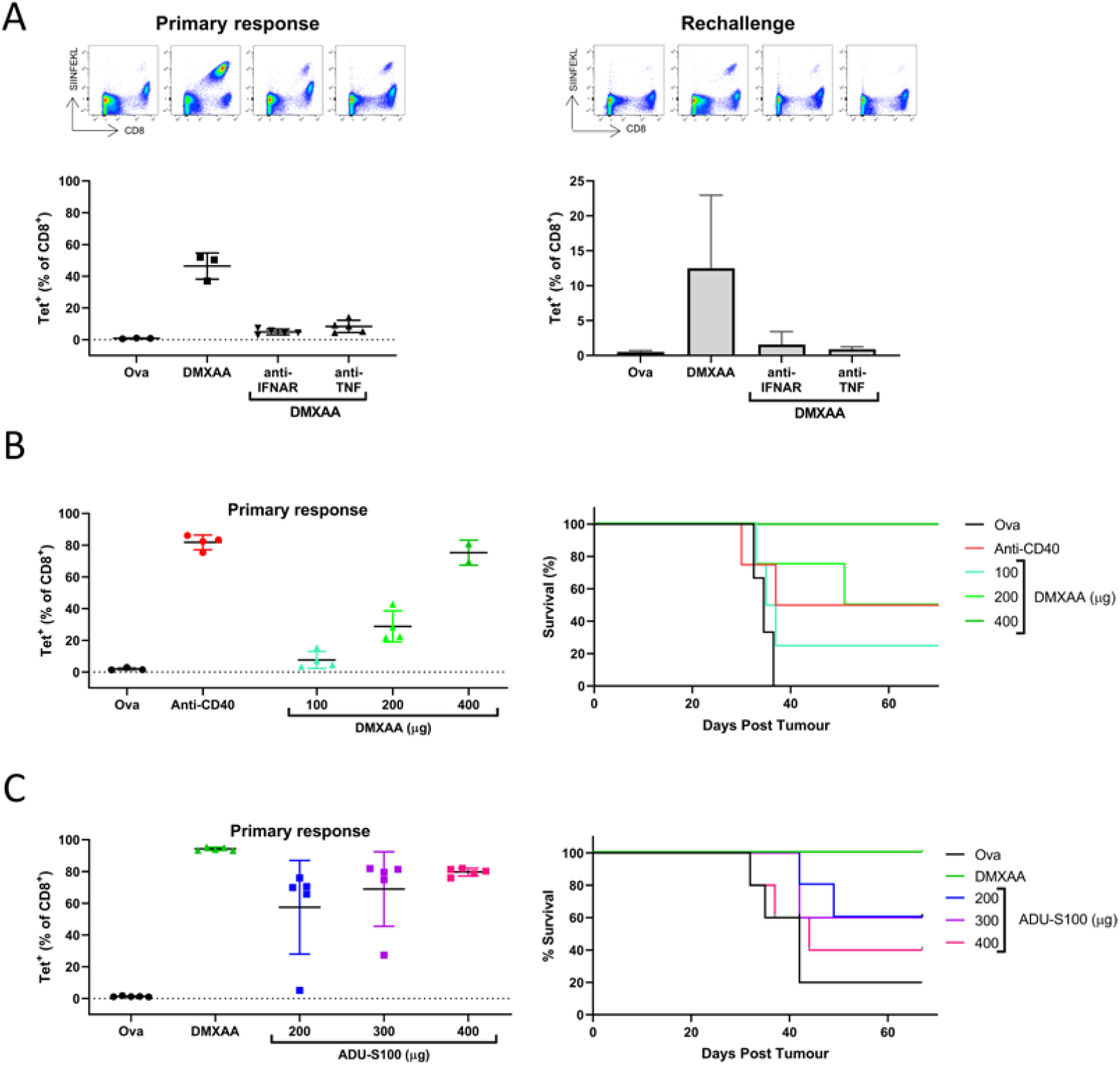
STING agonist mediated adjuvancy leads to a robust memory response and better tumour protection. Mice challenged as in Figure 1 were subject to rechallenge once primary CD8^+^Tet^+^ response had contracted to <10% of total CD8s. (**A**) Re-challenge of mice receiving DMXAA with and without IFNAR and TNF blockade: Left panel, primary response showing Tet^+^ cells in blood on day 7 peak; right panel, re-challenge with SIINFEKL showing Tet+ cells in blood at and day 7 peak of response. (**B**&**C**) E.G7 tumour challenge; primary response (left), post challenge survival plot (right). (**B**) Tumour control in mice dosed with varying dose of DMXAA. (**C**) Tumour control in mice receiving varying doses of ADU-S100. Each point represents an individual mouse, mean is indicated with error bars showing SD.

## Discussion

Innate immune adjuvants are sought for combination with approaches promoting adaptive immunity with the aim of enhancing immune responses and maximising long-term antigen specific memory. One of the most promising innate activators are STINGa. However, despite promising results in preclinical models, these innate activators are yet to translate successfully to the clinic. Better understanding of their mechanism of action and the properties affording good adjuvancy for adaptive immunity will aid clinical development.

Here we have demonstrated that synthetic STINGa are able to act as potent adjuvants to drive the expansion of antigen specific CD8s. In this context, STINGa outperform CD40 ligation with the contrasting effects induced likely reflect fundamental mechanistic differences inherent to their respective inductions of Type-I versus Type-II IFN.[3, 33]

We confirmed that synthetic STINGa are contingent on STING for their adjuvancy and that in this setting Type-I IFN and TNFα are required. We compared the performance of two distinct synthetic STINGa: the murine active xanthenone analogue, DMXAA and the clinically relevant cyclic dinucleotide, ADU-S100.[24-26] Both acted as effective adjuvants with the latter less potent in driving CD8 expansion, even at the maximally tolerated dose. The predicted ∼8 fold greater potency of ADU-S100 versus DMXAA for STING activity[37, 38] would suggest the inverse would be true. Previous studies with ADU-S100 have proposed a bell-shaped curve for dose response for infiltration of CD8^+^s to tumour upon intratumoural administration,[37] but we observed no deviation from a positive dose response for CD8 expansion up to and inclusive of the maximum tolerated dose.

DMXAA has a reported half-life range of 4-12 hours,[44] whereas, ADU-S100 is cleared more rapidly, within 10-23 minutes in plasma[45] and it is possible that adjuvancy in this setting is influenced by the balance between potency and half-life; DMXAA possessing lower potency but slower clearance providing more sustained STING activation. Although this represents an interesting explanation for their differential effects, the half-lives of both compounds are relatively short compared to anti-CD40 which can persist for more than a week,[46] therefore, it is likely that potency is probably more important.

A candidate mechanism by which STINGa may induce a greater antigen specific CD8 expansion compared to anti-CD40 is via modulation of the immune checkpoint PD-1. PD-1 is rapidly upregulated once T-cells are activated[39] and acts to inhibit further TCR signalling and suppress T cell activity.[40] Our data demonstrate that expression of PD-1 on expanding T cells during antigen challenge with STINGa treatment was significantly lower compared to anti-CD40, which may explain their enhanced capacity to proliferate and, in the case of DMXAA, dominate the T cell pool. These differences in PD-1 were observed in all monitored CD8 compartments (Tet^-^ and Tet^+^). The observed differential response in PD-1 induction between DMXAA and ADU-S100 also offers a potential explanation for the lower adjuvancy provided by ADU-S100, suggesting its activity is limited by its potency.

STING is expressed widely by immune cells, with high expression in lymphocytes and relatively low expression in myeloid populations in mice. Given this, we next sought to determine which cells played a role in the CD8 expansions induced by STINGa. Previous reports have shown that T cells are uniquely sensitive to STINGa-mediated apoptosis.[41] In agreement, we found that STINGa exposure significantly inhibited T cell proliferation during CD3 stimulation in vitro, which, coupled with data that showed adjuvancy was lost in STING KO recipients, supports our assertion that the effect is mediated by accessory cells and not through direct effects on CD8s.

Previous reports, including our own, have shown that STING activation can induce the reprogramming of macrophages to a proinflammatory (M1-like) phenotype and the maturation of antigen presenting cells.[16, 23] Therefore, we sought to explore the potential roles of monocytes and macrophages in the STINGa induced CD8 responses via the blockade of CSF1 signalling. Our data demonstrate that inhibiting this axis led to a clear abrogation of STINGa induced Tet+ expansion strongly supporting a role for inflammatory monocytes and/or macrophages in the CD8 expansion observed. These findings support the importance of accessory cells to provide a favourable cytokine milieu for the induction of effective T cell responses in this setting. The negative impact of STING agonism on T cells also highlights the potential importance and detrimental impact of potency on responses. The contention being that the drive for more and more potent agonists may in fact elicit detrimental impacts on immunostimulatory adjuvancy.

The most important characteristic of any vaccine response is the strength of memory evoked. CD8 T cell memory has recently been shown to arise from de-differentiation of memory precursor effector cells (MPECs).[42] The generally held consensus is that within an effector population there reside short lived effector cells (SLECs) and MPECs.[43] It is the MPECs within the effector population that are destined to de-differentiate into CM cells whilst SLECs do not persist.[47] The synthetic STINGa utilised here produced robust effector expansion during antigenic challenge, but also stimulated a greater number of CM cells compared to anti-CD40 that persisted and expanded throughout the response. The CM cells became the dominant population in STINGa dosed animals much earlier compared to those treated with anti-CD40. These effects may be due to the dominance of Type-I IFN as downstream effector of STING as opposed to Type-II IFN with anti-CD40.[3, 33] When comparing DMXAA with ADU-S100 it was clear that despite the smaller Tet+ expansion induced by the latter, both STINGa produced similar early effects on CM dominance, strongly suggesting this is a general property of STING activation and independent of the size of the initial CD8 T cell expansion.

Together, this supports the contention that within the effector population induced during STINGa treatment a larger proportion of MPECs are produced compared with anti-CD40; again perhaps as a result of Type-I versus II IFN. Irrespective of the molecular basis, it seems clear that STINGa induced EM populations more rapidly transition into CM cells leading to more robust antigen specific memory.

Recent evidence suggests that STING expression and activation within T cells supports their proliferation and longevity in response to antigen challenge in vivo.[48] Furthermore, Type-I IFN responses downstream of STING are required to confer a stemcell-like phenotype, supporting generation of CM cells through TCF1 expression. These findings provide a possible explanation for the improved memory response we demonstrate, although, the direct influence on T cells appears less important in our system than the role played by myeloid cells.

Our data implicate myeloid cells, likely monocytes and/or macrophages, in the improved memory induction by STINGa. The propensity for STINGa to activate macrophages towards a more M1-like phenotype may help promote T cell memory through an appropriate cytokine milieu during priming. One such mechanism might be the effect of Type-I IFN on expression of CXCR3 ligands and subsequent influence on cell fate.[49] It is thought that a stronger inflammatory environment during priming drives T cells towards an effector fate whereas in a less inflammatory context T cells tend towards a memory fate. This may be the key difference between the Type-II IFN response mediated by anti-CD40 and the synthetic STINGa inducing a Type-I IFN dominant response.

This effective and rapid induction of robust memory represents an important finding that demonstrates a significant advantage of STINGa over other adjuvant strategies. To our knowledge, this is the first demonstration of superior CM formation using synthetic STINGa as an adjuvant in a CD8 T cell vaccine model. We also showed that the memory pool generated in response to STINGa is capable of robust recall upon rechallenge with the SIINFEKL peptide. This recall was absent when Type-I IFN or TNF alpha blockade were present during the primary challenge. This is not surprising given the lack of a primary response under these conditions. The memory recall produced in DMXAA treated mice evoked superior protection from Ova expressing tumour compared to anti-CD40 despite a similar primary response, demonstrating the functional effect of the dominant CM population. Only the highest tolerated dose of DMXAA provided this improved protection, which agrees with our assertion that a balance of dosing with potency and half-life is required for STINGa to have maximum effect. Accordingly, ADU-S100 at the highest tolerated dose provided less protection compared to the highest dose of DMXAA, which may indicate a detrimental effect on T cells despite evidence of a similar memory response. It is well established that immunological memory is defined by the pool of antigen-specific cells whose frequency governs rapid response to pre-existing antigen[50] Therefore, in our tumour vaccine setting, it is plausible that higher level of protection against the tumour relates to higher proportion of memory cells generated following DMXAA administration than that with ADU-S100.

We have demonstrated that synthetic STINGa can be used as effective adjuvants to soluble antigen in a vaccine setting, providing robust CD8 T cell expansion dependent on Type-I IFN and TNFα. This expansion is characterised by reduced PD-1 induction on CD8s when compared with the immune modulating adjuvant anti-CD40. We also show that STINGa induce better memory responses, causing an earlier central memory dominance which leads to larger responses to secondary challenge in the forms of peptide and antigen expressing tumour, resulting in better tumour control and mouse survival. To our knowledge, this is the first demonstration of synthetic STINGa inducing long-term CD8 T cell memory through vaccination, with direct implications for tumour control.

## Declarations

### Ethics approval and consent to participate

Animal experiments were conducted under UK Home Office licence number P81E129B7 following review by the Scientific and Ethical Review Group and approval by the Home Office Animal Welfare Ethical Review Board (AWERB) at the University of Southampton

### Consent for publication

All authors have given their consent for the publication of the work enclosed

### Availability of data

All data relevant to the study are included in the article or uploaded as supplementary information.

### Author contributions

L.D.T., L.N.D., M.C.T., and K.L.C. performed experiments. L.D.T. and L.N.D. analyzed and interpreted data. T.I., M.M., S.R.W and C.R. provided key reagents. L.D.T., L.N.D., M.S.C. and S.A.B designed experiments. L.D.T., L.N.D., S.R.W., M.S.C. and S.A.B. wrote the manuscript. All authors contributed to manuscript revision and read and approved the submitted version.

## Acknowledgements

We are grateful to the staff of the University of Southampton Biomedical Research facility for their technical support. We thank John Cambier and Jan Rehwinkel for generously providing STING KO (*MPYS*^*-/-*^) mice. We also thank Leon Douglas and Patrick Duriez from the ECMC/CRUK Protein Core Facility for providing key reagents. We thank Nicola Wallis, Astex Pharmaceuticals, for her support with this project.

## Abbreviations

STING: Stimulator of interferon genes
IFN: interferon
TNF-α: tumour necrosis factor alpha
PRR: pattern recognition receptor
ER: endoplasmic reticulum
CDN: cyclic dinucleotides
cGAS: cyclic GMP-AMP synthase
TBK1: tank binding kinase 1
IRF3: interferon regulatory factor 3
IFNAR: interferon-alpha/beta receptor
DC: dendritic cell
TME: tumour microenvironment
STINGa: Sting agonist
DMXAA: 5,6-dimethylxanthenone-4-acetic acid
cGAMP: cyclic GMP-AMP
PD-1: Programme cell death receptor 1
Ova: ovalbumin
Tet: tetramer
CD8^+^: CD8 T cell

