## Supplementary Figures for "Synthetic STING agonists elicit powerful vaccine adjuvancy providing robust central memory and anti-tumour effects"

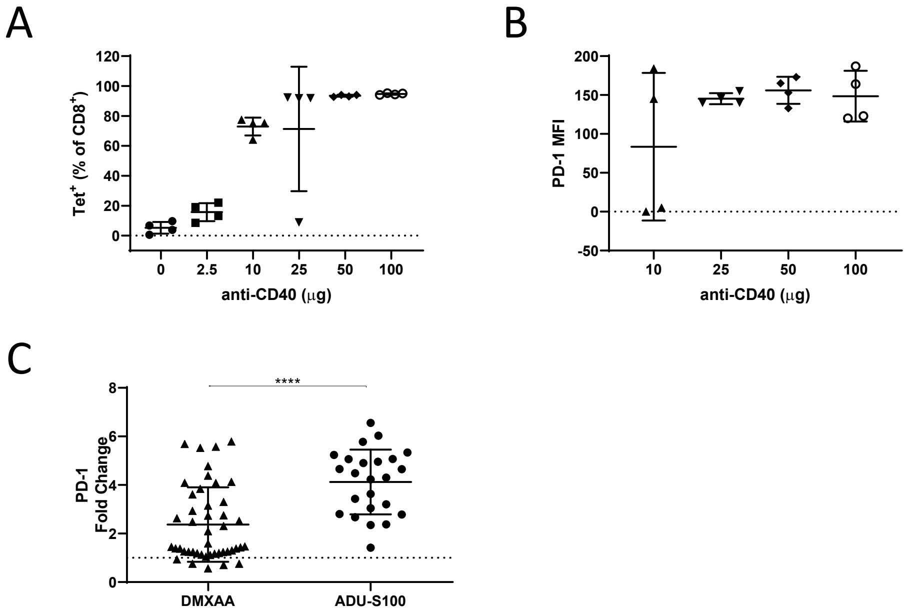


**Supplementary Figure 1**. **Anti-CD40 dose effect on Tet+ expansion and PD-1 induction; PD-1 levels differ between DMXAA and ADU-S100.** (**A**) Peak expansion of Tet+ cells and (**B**) levels of PD-1 expression, given by elevating doses of anti-CD40 (3/23). (C) Direct comparison of PD-1 induction by DMXAA and ADU-S100. Each point represents an individual mouse, mean is indicated with error bars showing SD. *P* values were determined using One-way ANOVA with Tukey's adjustment. ns = not significant; * *P*<0.05; ** *P*< 0.01; *** *P*< 0.001; *****P*<0.0001.

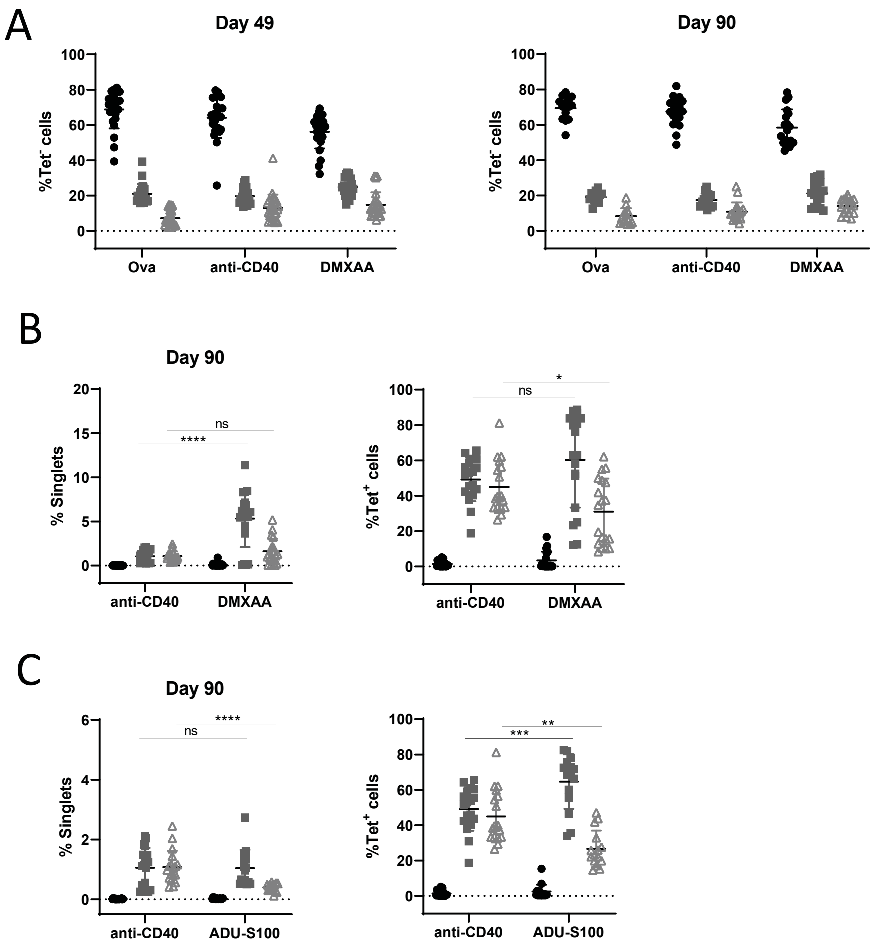


**S̲upplementary Figure 2. Long term memory phenotypes induced by STING agonists.** (**A**) Memory phenotypes of Tet$^-$ populations on day 49 and 90. Memory phenotypes of Tet$^+$ cells on day 90 induced by (**B**) DMXAA and (**C**) ADU-S100, left as %Singlets and right as %Tet$^+$ cells. Each point represents an individual mouse, mean is indicated with error bars showing SD. *P* values were determined using One-way ANOVA with Tukey's adjustment. ns = not significant; * $P$<0.05; ** $P$< 0.01; *** $P$< 0.001; ****$P$<0.0001.
